# Human brain activity during mental imagery exhibits signatures of inference in a hierarchical generative model

**DOI:** 10.1101/462226

**Authors:** Jesse Breedlove, Ghislain St-Yves, Cheryl Olman, Thomas Naselaris

## Abstract

Humans have long wondered about the function of mental imagery and its relationship to vision. Although visual representations are utilized during imagery, the computations they subserve are unclear. Building on a theory that treats vision as inference about the causes of sensory stimulation in an internal generative model, we propose that mental imagery is inference about the sensory consequences of predicted or remembered causes. The relation between these complementary inferences yields a relation between the brain activity patterns associated with imagery and vision. We show that this relation has the formal structure of an echo that makes encoding of imagined stimuli in low-level visual areas resemble the encoding of seen stimuli in higher areas. To test for evidence of this echo effect we developed imagery encoding models—a new tool for revealing how imagined stimuli are encoded in brain activity. We estimated imagery encoding models from brain activity measured with fMRI while human subjects imagined complex visual stimuli, and then compared these to visual encoding models estimated from a matched viewing experiment. Consistent with an echo effect, imagery encoding models in low-level visual areas exhibited decreased spatial frequency preference and larger, more foveal receptive fields, thus resembling visual encoding models in high-level visual areas where imagery and vision appeared to be almost interchangeable. Our findings support an interpretation of mental imagery as a predictive inference that is conditioned on activity in high-level visual cortex, and is related to vision through shared dependence on an internal model of the visual world.

---

Why do humans have mental images, and how do mental images differ from the ones we see? Two millennia of efforts to explain mental imagery have yielded wildly varying and inconsistent answers to these questions (*1, 2*). However, the relatively recent ability to measure activity in the brains of humans as they imagine has narrowed the range of possible answers around some key empirical facts: imagery engages the same brain areas as vision (*3, 4*), and activity in these areas encodes unambiguously visual representations (*5–8*). Nonetheless, the visual system appears to operate in a different regime during imagery, exhibiting altered (relative to vision) functional connectivity (*9, 10*), dynamics (*11*), and activity patterns (*12, 13*). A successful theory of mental imagery must thus explain what computations are subserved by visual representations that emerge independently of visual input, and are encoded in brain activity patterns that differ markedly from those observed during vision.

Here we formalize and test a theory of mental imagery that builds on an influential theory of vision (*14–17*). Under the theory, activity patterns in distinct visual areas, *r*_0_*, …, r_L_*, co-occur with the same probability as the features they encode co-occur in the real world (*18*). The joint distribution that specifies these probabilities *p*(*r*_0_*, …, r_L_*), is a hierarchical generative model (HGM), so-called because features at one stage of the hierarchy (say *r_L_*) are causally responsible for generating the features at lower stages (*r_l<L_*) in the way that, for example, the feature “sky” generates the feature “blue” (*14*). Seeing an image *s* corresponds to conditioning—or “clamping”—activity at the bottom of the hierarchy (e.g., the retina) so that *r*_0_ = *s* while the remaining activity patterns are sampled from a *posterior* distribution *p*(*r*_1_*, …, r_L_|r*_0_ = *s*). The average activity state for the *l*^th^ visual area

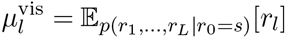

therefore encodes the expected *cause* of the retinal stimulus.

We treat mental imagery as a subtly but importantly different conditional inference within the same HGM. Specifically, we propose that imagining *s* corresponds to clamping activity in at least one visual area to the expected activity pattern evoked by seeing 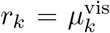, while clamping the retina to some uninformative value (e.g., *r*_0_ = 0). Activity in the visual system is then distributed according to the conditional probability distribution 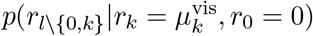. The average activity pattern for the *l*^th^ visual area

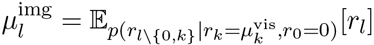

therefore encodes an expected *consequence* of the cause specified by clamping activity in a high-level visual area.

Treating mental imagery as an inference complementary to vision implies a compelling answer to the question of why humans have mental images: we have them because inferring the visual consequences of a predicted or remembered cause is useful during both deliberate cognition and ongoing perception of visual stimuli (*19, 20*). The particular form of inference we propose has face-validity, as it is an effective procedure for synthesizing images from abstract representations (*21*). Perhaps most importantly, it leads to testable predictions about how activity differs between imagery and vision. To make this clear we note that, in general, the expected activity pattern during vision at some higher stage *l* + *d* can be expressed as some (nonlinear) transformation 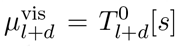 of the activity pattern at stage 0 (i.e., the stimulus *s*) to an activity pattern at *l* + *d*. Furthermore, in a strictly hierarchical architecture the transformation from any one stage to another can be decomposed into transformations between intervening stages (Fig. 1a), e.g., 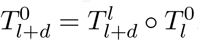. Since during imagery the dominant source of variance is the clamped layer, 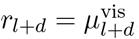, it follows that

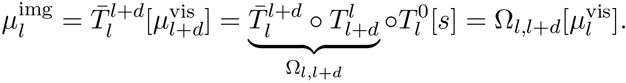

where *T*̄ transforms activity patterns from stage *l* + *d* to *l* (we use bar notation to indicate transforms from a higher to a lower stage). The mean activity pattern during imagery thus differs from the mean activity pattern during vision by a transformation, denoted Ω_*l,l*+*d*_, that is formally an echo. Under the echo transformation, an imagery activity pattern at any one stage will resemble a visual activity pattern that has been fed forward to the clamped stage, then fed back to the original stage. Unless this fed-back echo of the visual activity pattern is lossless, the encoding of imagined stimuli in the imagery activity pattern *beneath* the clamped stage will more closely resemble the encoding of seen stimuli *at* the clamped stage (*22*). This effect will appear strongest in the stage most distant from the clamped one, because it is there that the encoding of seen stimuli will be most different from encoding at the clamped stage. These echo effects are the titular signatures we expect to observe if mental imagery and vision correspond to the distinct forms of inference we hypothesize.

**Figure 1:**
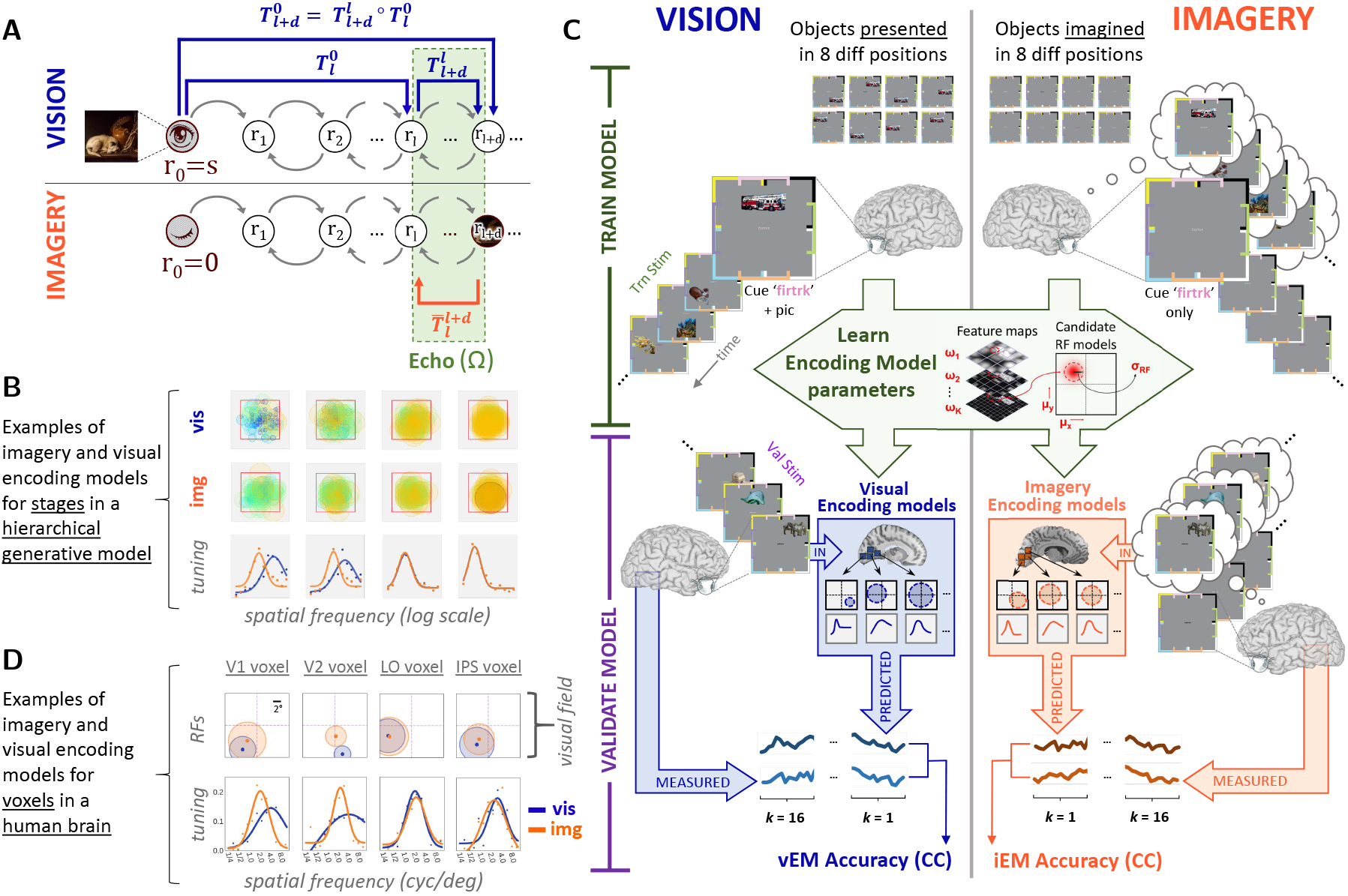
Characterizing differences in the encoding of seen and imagined stimuli with visual and imagery encoding models (**A**) Activity patterns in a hierarchical generative model (HGM). During vision the visual activity pattern at a processing stage, *r_l_*_+*d*_, is determined by a transformation 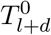 (long blue arrow) of activity (denoted *s* for stimulus) at the sensor stage 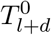 is equivalent to a composition of transformations (shorter blue arrows) of activity patterns between intervening stages. During imagery *s* = 0 but at least one processing stage (*r_l_*_+*d*_ in this example) is clamped to its visual activity pattern. Imagery activity patterns beneath the clamped stage (e.g., *r_l_*) differ from their visual activity patterns by an echo Ω. The echo is a transformation from the current to the clamped layer (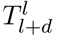, shortest blue arrow) and from the clamped layer back (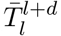, orange arrow). (**B**) The echo effect in an HGM. Receptive fields (top panels) and spatial frequency tuning functions (bottom) for vision and imagery are identical near the clamped stage (far right panel) but diverge beneath it (left panels). (**C**) Data and procedures for estimating visual (left, vEM) and imagery (right, iEM) encoding models. Whole-brain fMRI (7T) measured BOLD activity as subjects viewed or imagined 64 unique stimuli at 8 distinct locations (left). The color of the six-letter cue for each stimulus coded a location bounded by a visible bracket. Model estimation (center) was applied separately to visual and imagery data, resulting in a distinct vEM and iEM for each voxel. Model prediction accuracy was k-fold cross-validated by computing Pearson correlation between predicted and measured activities on held-out data. (**D**) iEM (orange) and vEM (blue) for single voxels that exemplify population-level trends in visual area V1, V2, lateral occipital cortex (LO) and intraparietal sulcus (IPS).

Note that expressions for both the visual and imagery activity patterns define two distinct encoding models (*23*), 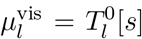 and 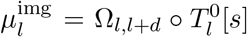, that differ only by an echo. It should therefore be possible to test for an echo effect on activity patterns in the human brain by estimating and then comparing visual and imagery encoding models (note that these expressions direct us to train and test each type of model using the same visual stimuli). If an echo effect holds then tuning to imagined features revealed by imagery encoding models in lower visual areas should be significantly different from the tuning to seen features revealed by visual encoding models. In particular, tuning to imagined features in lower visual areas should more closely resemble tuning to *seen* features in higher areas.

To illustrate the echo effect we compared visual and imagery encoding models estimated from the activities of units in an HGM (*21*) as it performed the distinct conditional inferences corresponding to vision and mental imagery. By design, the tuning of units to basic visual features varied across the model hierarchy in a manner consistent with known variation in tuning across the hierarchy of areas in human visual cortex. Thus, during vision units at higher stages exhibited lower spatial frequency preference, and larger receptive fields with correspondingly greater foveal coverage than units at lower stages (Fig. 1B,S4A, top panels). In contrast, during imagery units below the clamped stage exhibited— relative to vision—reductions in spatial frequency preference, increases in receptive field size and shifts of receptive field center toward the fovea (Figs. 1B, S4D). Thus tuning to imagined features at lower stages more closely resembled tuning to seen features at the clamped stage (Fig. 1B, bottom). The size of these changes increased with distance from the clamped stage (Fig. S4).

We conducted an fMRI experiment to determine if these and other signatures of inference in an HGM could be observed during mental imagery in the human brain (Fig. 1C). We measured whole-brain BOLD activity as participants viewed and then in separate sessions were cued to imagine previously memorized pictures of objects at different positions in the visual field. Distinct voxelwise visual and imagery encoding models (*24*) were estimated from activity measured during viewing and imagery sessions, respectively (Fig. 1C). Encoding models specified tuning to spatial frequency and a receptive field location and size for each voxel. Any voxel sensitive to the cues was discarded.

Predictions made by imagery encoding models exceeded a threshold on accuracy for many voxels (subsequent analyses refer to above-threshold voxels only) in all visual cortical areas considered here (Fig. 2A–C). Importantly, predictions of the imagery encoding models were accurate enough to identify the position of (Fig. 2D) and object in (Fig. 2E) the imagined stimuli (*5, 25*). Accurate identification of imagined objects would not be possible if variation in spatial attention, eye position or visual cues were the determinants of prediction accuracy. These results thus license us to inspect the parameters of the imagery encoding models for differences in the encoding of imagined and seen stimuli.

**Figure 2:**
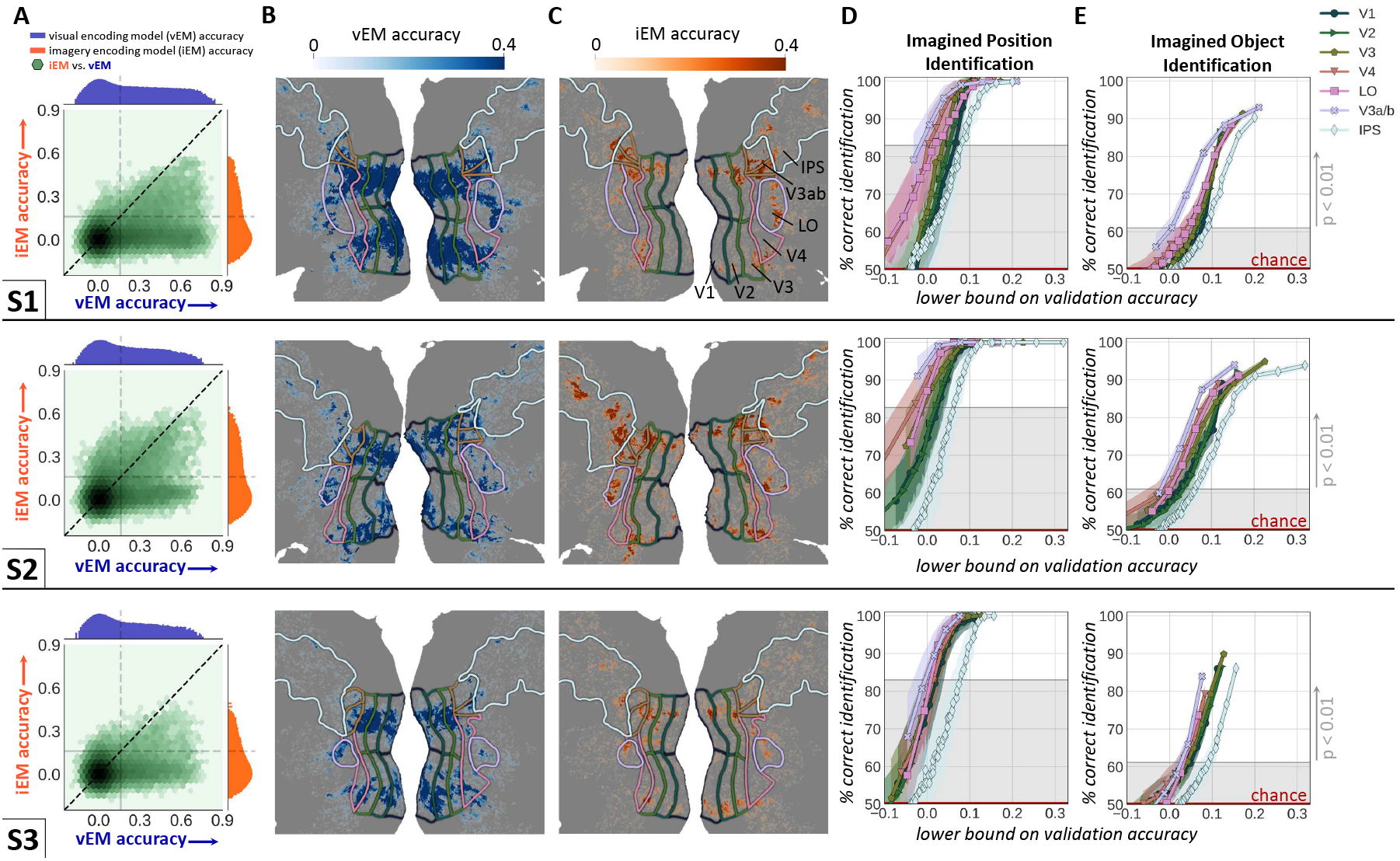
Validation of imagery encoding models. (**A**) Joint histogram (green) and marginal histogram of prediction accuracy for imagery (orange) and visual (blue) encoding models across all voxels for subjects 1-3. The imagery encoding model (iEM) makes accurate predictions of imagery activity (Pearson correlation ≥ 0.16, *p* < .01; dashed grey lines) in all subjects. (**B**) Prediction accuracy (colorbar) of the visual encoding model (vEM) mapped on the flattened cortical surface. (**C**) iEM prediction accuracy. (**D**) Imagined location identification. Curves show percentage of correct pairwise identification (colored shading indicates ± SE; gray shading indicates statistical significance threshold of *p* < .01 (permutation test)) for subpopulations of 500 voxels in visual area. Ordering along x-axis is by lowest prediction accuracy of all voxels in each subpopulation. (**E**) Imagined object identification. Format as in D.

If an echo effect were induced by clamping in a high-level visual area, we should expect prediction accuracy of imagery and visual encoding models to be close to parity in this area. This was true in intraparietal sulcus (IPS), a collection of visual areas at the highest stage of processing considered here (Fig. 3A,B). Relative prediction accuracy of the imagery encoding model decreased with descent toward primary visual cortex (V1). This gradient in relative encoding model prediction accuracy is most likely due to a matched gradient in relative signal-to-noise (SNR; Fig. 3C). Interestingly, our theory predicts this gradient in relative SNR, and that the loss of SNR during imagery depends only on a reduction in signal amplitude, since clamping at an additional stage during imagery will *reduce* noise during imagery relative to vision (Fig. S3). Distinct signal (Fig. 3D) and noise (Fig. 3E) measures were consistent with this prediction.

**Figure 3:**
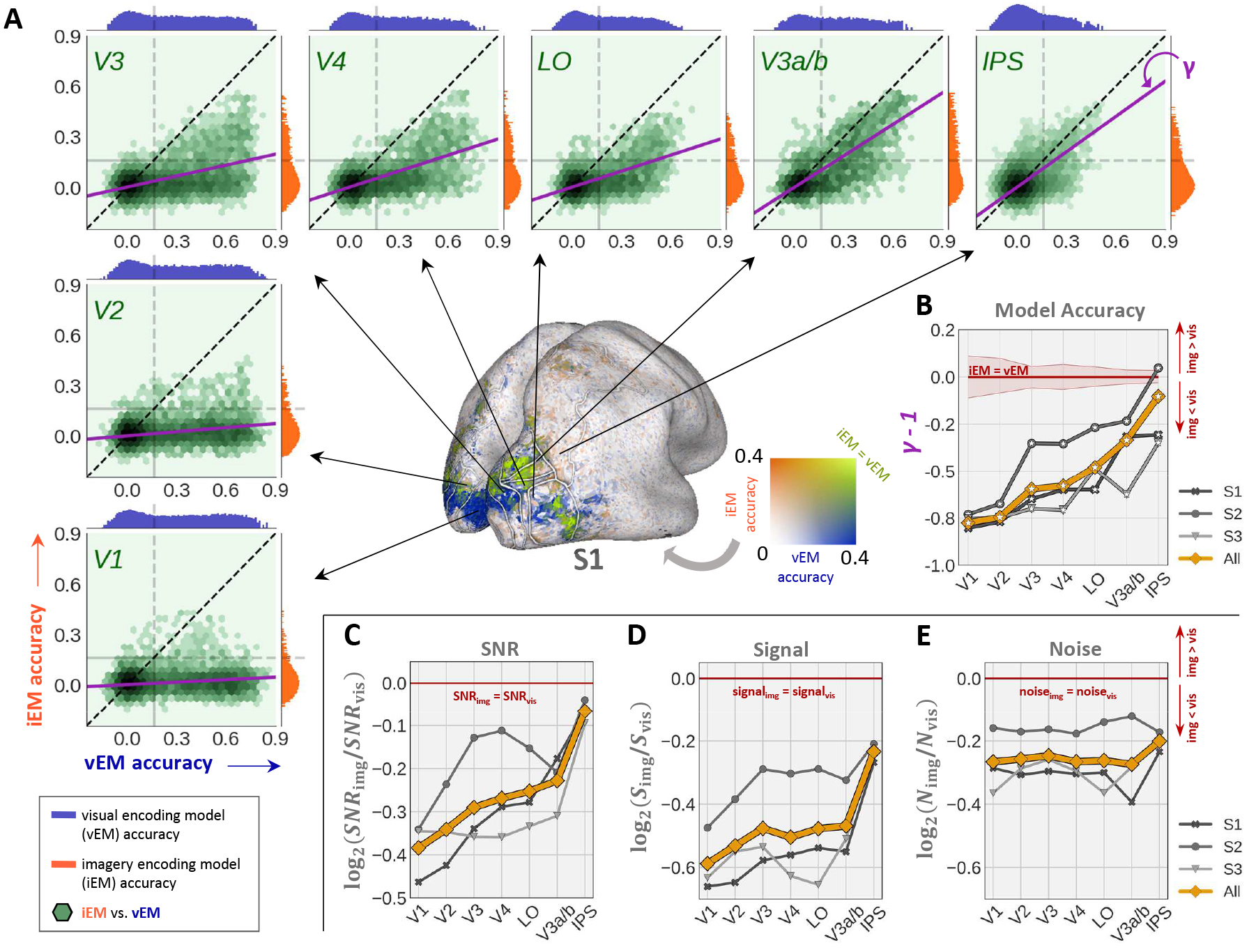
Relative prediction accuracy of imagery encoding models (iEM) across visual areas (**A**) Joint and marginal prediction accuracy histograms (format as in 2A) for indicated area (subject 1 only; ordering of visual areas follows (*34*)). Purple line shows slope (*γ*) of best linear fit of iEM to vEM prediction accuracy. Inflated surface map shows relative prediction accuracy (2d colormap) of the iEM and vEM. (**B**) Difference from parity (*γ* 1) for each area. (**C**) Median signal-to-noise ratio (SNR) for imagery activity relative to visual activity. (**D**) Relative activation amplitude (*S*) and activation noise (*N*).

When an echo is induced by clamping at a high-level visual area, imagery spatial frequency preference during imagery should decrease relative to visual spatial frequency preference with descent toward V1 (Fig. 1B bottom panels, S4D). Imagery spatial frequency tuning peaks were consistent with this echo effect. Unlike encoding model prediction accuracy, loss of SNR in early visual areas cannot account for these effects (Fig. S11).

Another effect of an echo induced by high-level clamping is that imagery receptive field sizes should move (relative to vision) toward the fovea and dilate with descent toward V1 (Fig. 1B, top two panels, S4A–C). Although receptive field attributes were highly variable across subjects these position (Fig. 4F–H) and size (Fig. 4I) effects were observed in combined subject data. Imagery receptive fields were relatively more foveal and larger in V1 for each subject.

**Figure 4:**
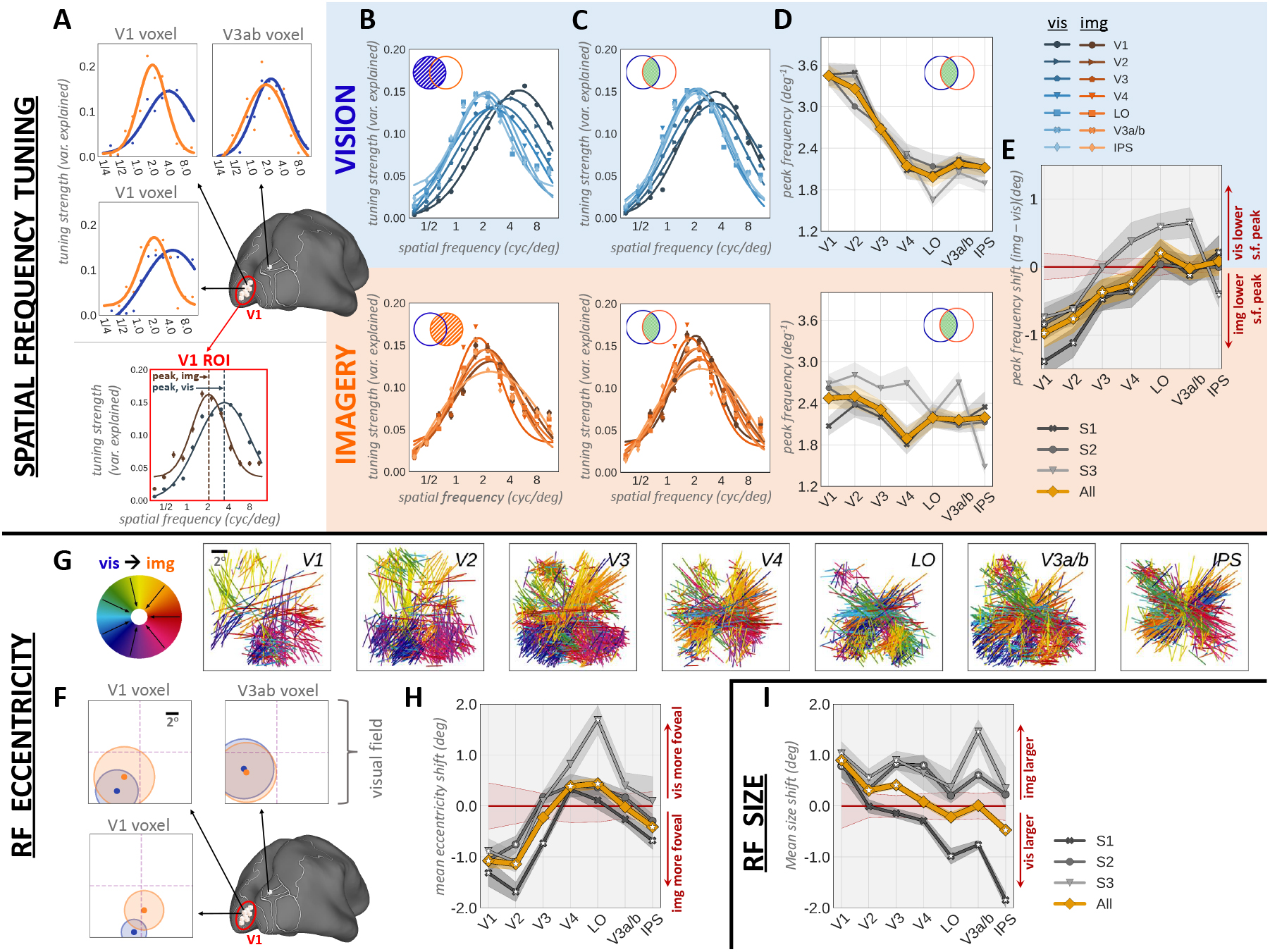
Differences between spatial frequency, receptive field location and size during vision and imagery. (**A**) Visual (blue) and imagery (orange) spatial frequency tuning curves for single voxels and population tuning curves (bottom) for V1. (**B**) Top: Population tuning curves during vision for all voxels in the indicated area that have an accurate vEM. Bottom: Population tuning curves during imagery for voxels that have an accurate iEM. Populations in top (blue circle in Venn diagram) and bottom (orange circle) plots are overlapping but not identical. (**C**) Population tuning curves for all voxels in the indicated area that have an accurate vEM and iEM. All subsequent panels use this population. (**D**) Peak spatial frequency. (**E**) Difference between peak spatial frequency during imagery and vision. (**F**) Example visual and imagery receptive fields (RF) (**G**) Orientation and magnitude (line segments) and direction (colorwheel at far left) of RF location shifts (same voxels as in D,E) from vision to imagery. (**H**) Average signed magnitude of shift in RF location from vision to imagery. Negative values indicate a shift toward fovea. (**I**) Average signed change in RF size from vision to imagery. Positive (negative) values indicate dilation (shrinkage). The red shaded area in E, H and I indicates significance level *p* < .01 (permutation test) for combined subject data (orange curve). In all panels asterix indicates significant difference from null value (red line, *p* < .01, permutation test; red shading indicates significance threshold for combined data); shading on curves indicates *±* SE.

The observed differences between visual and imagery activation amplitudes, noise levels, spatial frequency preference, receptive field location and size offer support for our formulation of mental imagery as inference in a generative model. To test this formulation we have introduced clamping (i.e., conditional inference in a generative model) as a potential mechanism by which high level visual areas may exert control over areas lower in the visual hierarchy. Our results suggest that in these experiments clamping occurred at least as high as V4; at and above V4 the effects of mental imagery on receptive field and tuning properties were much weaker than in areas lower on the hierarchy.

High-level clamping may explain why mental images lack the specificity of seen ones. High-level areas provide a poor substitute for the visual detail supplied by the retina during vision. This places an upper bound on the specificity of mental images that is most clearly revealed, in our analysis, by the reduced spatial frequency preference observed in lower areas during imagery. Our results thus suggest that limits to the specificity of mental images are built into the representational hierarchy of the visual system, and should remain even if the memories that specify mental images are perfectly encoded and recalled.

It is unclear how or if clamping relates to other forms of visual cognition, such as attention, that are difficult to cleanly disentangle from mental imagery. However, we note that across all subjects and analyses, the effects we attribute to clamping are strongest in lower areas, where the effects commonly attributed to attention are weakest (*26–28*). Interestingly, although the the differences between imagery and visual encoding models are small at and above V4, they do not entirely vanish; it is possible the subtle but significant differences in high-level areas reflect mechanisms related to attention, which are known to affect receptive field location and size in areas such as V3A/B (*29*) and those within IPS (*30, 31*).

By tying mental imagery to inference we have provided a potential explanation for how mental imagery could utilize visual representations but encode them in qualitatively different activity patterns. We have also given empirical support to the intuition that we imagine to “see” the visual consequences of visual predictions and memories. Our work also extends the power and relevance of the generative perspective on vision: while previous results relating vision to posterior inference have supplied evidence that representations in biological visual systems are adapted to the structure of the visual environment (*18, 32, 33*), the current results provide new evidence that these representations, once acquired, can emerge independently of retinal input (*18*), allowing the visual system to reason coherently about the visual environment even when there is nothing to see.

## Acknowledgments

We thank Dr. Truman Brown, Dr. Kendrick Kay, and Logan T. Dowdle for helpful comments and discussions on the manuscript. This work was supported by grant NIH R01 EY023384 to TN. CO, JB and TN designed the experiment. CO, JB and TN acquired the data. GSY and TN developed the theory. JB, GSY and TN analyzed the data. All authors contributed to writing and editing the manuscript.

